# Dynamic Changes in Lymphocyte Populations Establish Zebrafish as a Thymic Involution Model

**DOI:** 10.1101/2023.07.25.550519

**Authors:** Ameera Hasan, Jose J. Macias, Brashé Wood, Megan Malone-Perez, Gilseung Park, Clay A. Foster, J. Kimble Frazer

**Author notes:** Corresponding Author: J. Kimble Frazer. Email Addresses of Co-authors: Ameera Hasan, Jose J Macias, Brashe’ Wood, Megan Malone-Perez, Gilseung Park, Clay A. Foster, J. Kimble Frazer.

## Abstract

The thymus is the site of T lymphocyte development and T cell education to recognize foreign, but not self, antigens. B cells also reside and develop in the thymus, although their functions are less clear. During ‘thymic involution,’ a process of lymphoid atrophy and adipose replacement linked to sexual maturation, thymocytes decline. However, thymic B cells decrease far less than T cells, such that B cells comprise ∼1% of human neonatal thymocytes, but up to ∼10% in adults. All jawed vertebrates possess a thymus, and we and others have shown zebrafish (*Danio rerio*) also have thymic B cells. Here, we investigated the precise identities of zebrafish thymic T and B cells and how they change with involution. We assessed the timing and specific details of zebrafish thymic involution using multiple lymphocyte-specific, fluorophore-labeled transgenic lines, quantifying the changes in thymic T- and B-lymphocytes pre- vs. post-involution. Our results prove that, as in humans, zebrafish thymic B cells increase relative to T cells post-involution. We also performed RNA sequencing (RNA-seq) on *D. rerio* thymic and marrow lymphocytes of four novel double-transgenic lines, identifying distinct populations of immature T and B cells. Collectively, this is the first comprehensive analysis of zebrafish thymic involution, demonstrating its similarity to human involution, and establishing the highly genetically- manipulatable zebrafish model as a template for involution studies.

## Introduction

The thymus is a primary lymphoid organ whose specialized microenvironment fosters the development and selection of T lymphocytes, crucial to vertebrate immune function (1). Thymic involution refers to atrophy of, and declining T cell production by, the thymus with aging. This process begins prior to puberty, accelerates with sexual maturation, and continues in adulthood, resulting in diminished thymic epithelial space for T cells development (2). Thymic involution, and immunosenescence in general, are conserved in vertebrates, although the timing and rate of these processes vary by species (3, 4). Thymic involution has been studied in humans and mammalian models, but in other vertebrates, including zebrafish, involution is largely uncharacterized.

During involution, the thymic epithelial cells (TEC) that promote T cell development and selection diminish, leading to less naive T cell production (4–7). With fewer new T cells, T cell receptor (TCR) diversity also decreases, resulting in declining T cell function. (7–10). TEC reduction coincides with thymic adipocyte accumulation. These cells occupy non-epithelial thymic spaces and ‘infiltrate’ intra-thymic niches. (11) Contemporaneously, during involution, thymic perivascular spaces (PVS; non-thymopoietic regions that flank blood vessels) expand (12). Recently, the thymic PVS was recognized as a thymic plasma cell (PC) reservoir (13). Although often considered a T cell organ, the thymus also contains B lymphocytes with unique phenotypes compared to B cells elsewhere, (14, 15) with some thymic B cells participating in T cell development and selection (16). In humans, unlike T cells, thymic B cells do not decline precipitously with involution. In fact, they actually rise (relative to T cells) from ∼1% of total thymocytes pre-involution to ∼10% post-involution (13).

Although some aspects of involution vary by species, maximal thymic involution generally coincides with puberty (17–20). In teleost (bony) fish, involution exhibits variable timing, extent, and permanent vs. seasonal occurrence, suggesting environmental and genetic factors regulate involution (17). Despite this, thymic development and anatomy are similar between fish and humans (21). Recent single-cell RNA sequencing (scRNA-seq) results described the zebrafish thymus transcriptional landscape and demonstrated the presence of pre-B cells in zebrafish (22). *D. rerio* are widely-used in developmental studies due to their early life stages being transparent-to-translucent, their small size, and their rapid development—features ideal for live imaging (23). Paired with their genetic tractability, zebrafish have enhanced our understanding of thymopoiesis. (24–27). However, because involution occurs later, in older non-transparent/translucent zebrafish, much about *D. rerio* involution remains unknown.

Here, we studied several transgenic lines with fluorophore-labeled lymphocytes, at three timepoints, to comprehensively analyze zebrafish thymic involution. Specifically, we quantify the dynamic changes in thymic T and B cells during involution, accompanied by analysis of morphologic thymic structural changes. We also used double-transgenic fish to identify distinct thymic T and B cell sub-populations via RNA-seq. Our results establish *D. rerio* as a highly genetically-manipulatable model to learn the pathways and mechanisms governing the conserved vertebrate thymic involution process.

## Materials and Methods

### Zebrafish husbandry and transgenic lines

Zebrafish care was provided as previously reported (28, 29). Fish were housed in an aquatic colony at 28.5°C on a 14:10 hour light:dark circadian cycle. Experiments were performed according to protocols approved by the University of Oklahoma Health Sciences Center IACUC. For procedures, fish were anesthetized with 0.02% tricaine methanesulfonate (MS-222). The following transgenic lines were used: *lck:GFP* (27), *lck:mCherry* (gift from Aya Tal Ludin, Zon laboratory, Harvard University)*, cd79a*:*GFP*, *cd79b*:*GFP* (30), and *rag2*:*RFP* (31). Double-transgenic fish were made by breeding *cd79a*:*GFP* or *cd79b*:*GFP* transgenics to *rag2*:*RFP* and *lck:mCherry* transgenics.

### Fluorescent microscopy

Anesthetized fish were screened using a Nikon AZ100 fluorescent microscope. High exposure (1.5 s, 3.4× gain) settings were used to obtain images with a Nikon DS-Qi1MC camera. Images were processed using Nikon NIS Elements Version 4.13 software.

### Fish fixation, paraffin-embedding, sectioning, and H&E staining

Zebrafish at 3, 6, and 12 m were fixed in 10% neutral buffered formalin v/v for 2-3 days at room temperature. After fixation, samples were washed with 1% PBS three times and decalcified overnight in EDTA/Sucrose. Samples were then transferred to 70% ethanol, paraffin-embedded, and 60 to 120 sagittal sections cut (every 4µm), beginning at the eye surface. Every 5^th^ slide was H&E stained (20 µm apart). Using ImageJ^®^, thymic areas were measured for each H&E image, allowing thymic area and volume assessments.

### Flow cytometric and Fluorescence-Activated Cell Sorting (FACS) analyses

As previously described (29), thymi and marrow were dissected and placed in ∼500µl cell media (RPMI + 1%FBS + 1% Pen/Strep). Single cell suspensions were prepared by dissociating tissues with a pestle and passed through 35μm filters. GFP^hi^, GFP^lo^, and/or RFP^+^, mCherry^+^ cells were quantified and/or sorted from lymphoid and precursor gates using a BD-FACSJazz Instrument (Becton Dickinson, San Jose, CA, USA). Flow cytometric analyses were performed using FlowJo software (Ashland, OR, USA).

### 3D Model construction of zebrafish thymus

Aperio ImageScope .svs files were pre-processed, cropped, down-sampled, and exported into QuPath (32). Resulting .tif files were imported into Fiji/ImageJ (33) as virtual stacks and opened with the TrakEM2 plugin (34). Serial sections were aligned programmatically using Least Squares alignment (affine/rigid transform) and edited manually using anatomic landmarks. Layers were again aligned programmatically using Elastic alignment. Next, thymi were annotated manually using the brush tool and assigned to area_list objects. 3D models were generated from resulting thymic sections (downsample=40). Models were exported as binary .stl files and opened in MeshLab (35) for additional processing. Non-manifold edges and vertices were removed and resulting holes closed. Close vertices were merged and an HC Laplacian Smooth filter was applied 4x. Resulting models were then imported into Blender to create animations.

### 3’end RNA Sequencing

Using the fluorophore markers listed above, single- and double-positive cells were FAC-sorted from zebrafish thymus and marrow. Total RNA extraction was performed using a Promega Reliprep RNA extraction kit according to manufacturer’s instructions, generating ∼2.5 ug of RNA/sample. RNA concentrations were ascertained by fluorometric analysis on a Thermo Fisher Qubit fluorometer. RNA qualities were verified by Agilent Tapestation. Following QC, library construction was performed using the strand-specific QuantSeq 3’ mRNA-Seq Library Prep Kit FWD from Lexogen^®^ per manufacturer instructions. Briefly, 1^st^-strand cDNA was generated using 5’-tagged poly-T oligomer primers. After RNase digestion, 2^nd^-strand cDNA was generated using 5’-tagged random primers. Subsequent PCR with additional primers added complete adapter sequences with unique indices to demultiplex samples to initial 5’ tags, and amplified the library. Final libraries were assayed by Agilent Tapestation for size and quantity. Libraries were then pooled in equimolar amounts as ascertained by fluorometric analyses. Final pools were quantified by qPCR on a Roche LightCycler 480 instrument with Kapa Biosystems Illumina Library Quantification reagents. Sequencing was performed using an Illumina NovaSeq 6000, to a minimum depth of 20 million single-end 150bp reads/sample.

### RNA-Seq analysis

Data quality was assessed using FastQC (v.0.11.8). BBDuk from the BBMap suite of tools (v.38.22) (36, 37) was used for adapter and soft-quality trimming per manufacturer (Lexogen) recommendations and to remove rRNA mapping reads. Trimmed .fastq files were mapped to the GRCz11 genome using STAR with average unique mapping rates of 70-80%, resulting in a minimum of ∼10 million usable reads/sample. The *D. rerio* Ensembl v.92 transcriptome was used for gene annotation. Picard (v.2.18.14) and Qualimap (v.2.2.2-dev) (38) were used to assess alignment quality. FeatureCounts from the Rsubread (39) package (v.2.12.2) was used to generate gene counts using an alignment quality threshold of ≤10. DESeq2 (v.1.38.3) (40) was used for downstream processing and DE testing. Unique up-regulated markers were generated for each group in each lineage by comparing the fore-group of interest against the average of all other groups in the same lineage. Unless otherwise stated, target *p*-values and FDR thresholds were 0.05, with minimum absolute fold-changes of 1.5 (the lfcShrink function was used to predict fold-changes using the adaptive shrinkage estimator method) (41). A post-hoc filter was applied to marker genes to ensure each group of interest had at least 10 normalized counts in ≥ 2/3 of samples. All markers are distinct relative to other groups in the same lineage (B or T), but not necessarily to the entire dataset.

This dataset is deposited at NCBI GEO (https://www.ncbi.nlm.nih.gov/geo/) under accession # GSE237139; Reviewers, use this token to access: **udchcmckvtkxvgp**.

## Results

### Thymic Involution Causes Declining Fluorescence in Transgenic Zebrafish

Zebrafish thymic ontogeny has been thoroughly studied in embryos, but little is known about thymic involution in adults (4, 21). *D. rerio* thymic area increases during the first 8 weeks post-fertilization (wpf) and then declines with the onset of sexual maturity (18-52 wpf) (20). This thymic regression phenomenon, known as involution, occurs across vertebrates (3). To study the cellular and morphologic features of adult zebrafish thymi and changes linked with thymic involution, we performed a series of studies utilizing wild-type (WT) zebrafish and multiple transgenic lines. We first conducted serial fluorescent microscopy. Our prior work proved transgenic *lck:eGFP* (27) differentially labels T- and B-lineage cells in non-diseased and acute lymphoblastic leukemia (ALL)- prone zebrafish (28, 29). Similar *lck:mCherry* transgenic fish (gift from Aya Ludin, Zon laboratory, Harvard University) have not been reported upon previously. A third reporter line, *rag2:RFP*, labels immature lymphocytes of both the T and B lineages (42). We have also used two newer B cell-specific transgenic reporter lines, *cd79a:GFP* and *cd79b:GFP* (30), to analyze B cells and B-ALL (43). Using these five lines, we investigated gross thymic appearance over the first year of life.

By fluorescent microscopy, 3 month (m) fish of four lines exhibited robust thymic signals (Figure 1A, top row). These four lines (*lck:eGFP*, *lck:mCherry*, *rag2:RFP*, and *cd79a:GFP*) label various T cell populations (Park *et al.*, submitted; see attached pdf). In contrast, *cd79b:GFP* fish show higher B cell-specificity, explaining their weaker thymic fluorescence (Figure 1A, top right image). At 6m, thymic fluorescence was markedly lower in all five lines, with only *rag2:RFP* and *cd79a:GFP* fish retaining appreciable signal (Figure 1A, 2^nd^ row). By 12m, thymic signal was barely observable in all lines (Figure 1A, 3^rd^ row). These data, together with others’ prior work (3, 4, 20, 21) suggest the zebrafish thymic involution timeline depicted in Figure 1B. Notably, the 3m-6m “involution window” coincides with *D. rerio* attaining sexual maturity. To test this hypothetical window, we next performed several more rigorous studies to quantitatively and comprehensively define thymic involution in zebrafish.

**Figure 1:**
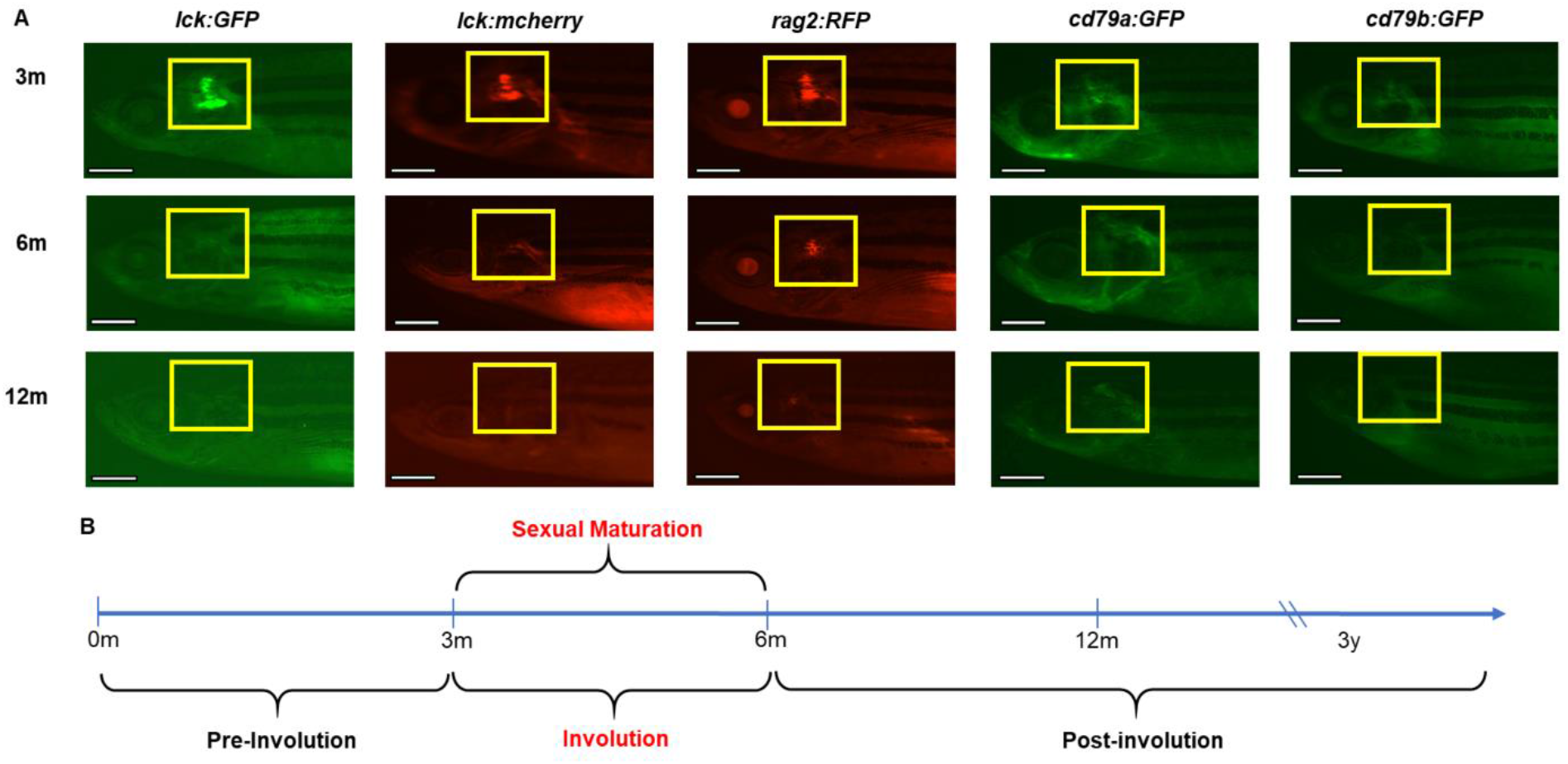
Declining thymic fluorescence in transgenic zebrafish during involution. (**A**) Fluorescent microscopy of transgenic lines with lymphocyte-specific fluorophores at 3, 6, and 12m. Yellow boxes denote thymic regions. Scale bars = 1mm. (**B**) Timeline of thymic involution, which peaks in the 3-6m window, coinciding with sexual maturity onset.

### Thymocyte Decline Explains Diminished Fluorescence During Involution

Imaging transgenic reporter lines can generally assess involution, but is decidedly imprecise. Therefore, to quantitatively measure changes in T and B cells during involution, we used flow cytometry to analyze thymocytes in *lck:eGFP, cd79a:GFP*, and *cd79b:GFP* fish at 3, 6, and 12m (Figures 2-4).

**Figure 2:**
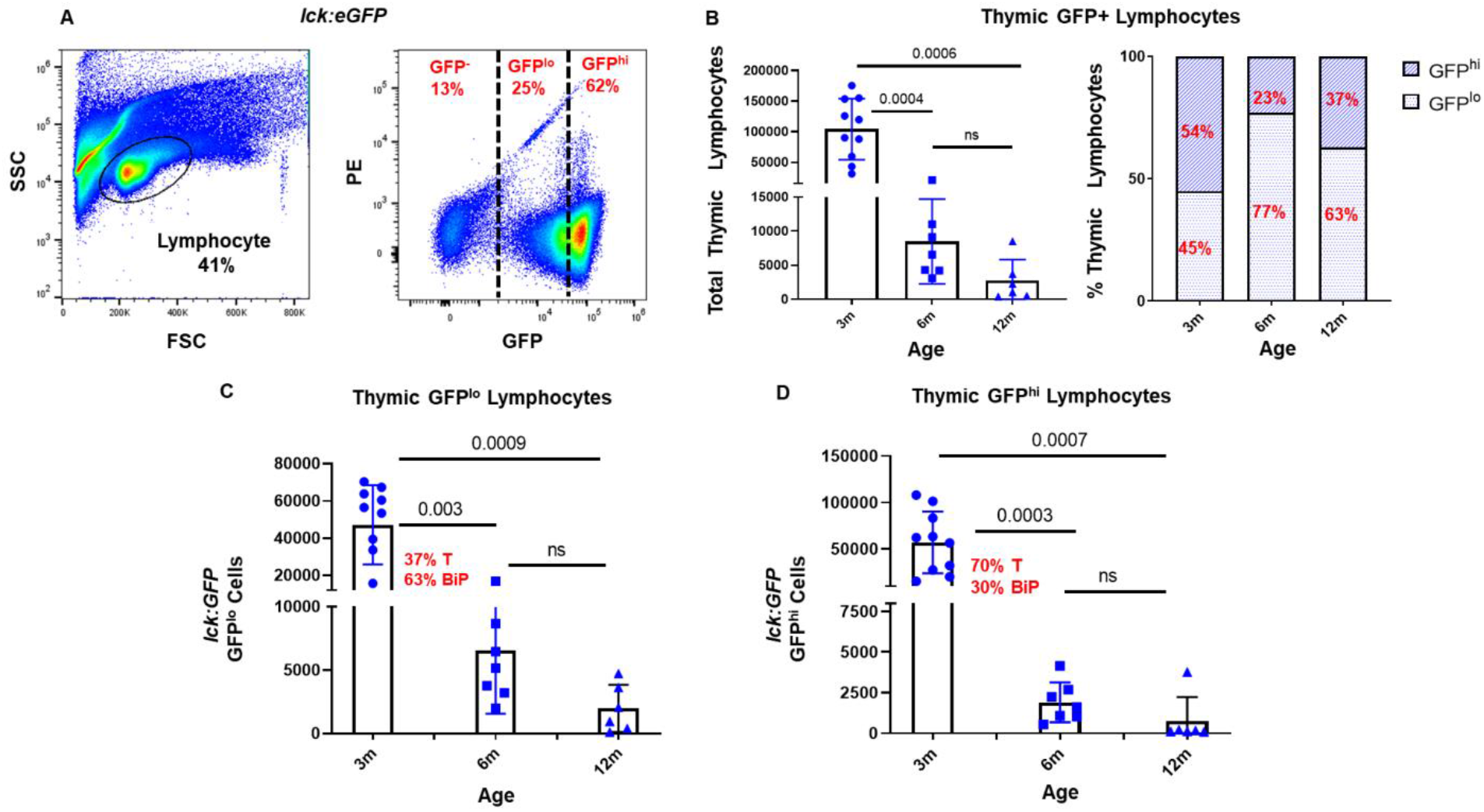
Quantification of thymocytes in *lck:eGFP* zebrafish during involution. (**A**) Sample thymus flow cytometry plots showing FSC/SSC-defined lymphoid gate and GFP^-^, GFP^lo^, and GFP^hi^ populations from this gate. (**B**) Quantification of total thymocytes (**left**) and % of GFP^lo^ vs. GFP^hi^ populations (**right**) **at** 3 (n=12 fish), 6 (n=7), and 12m (n=6). Declining (**C**) GFP^lo^ (37% T, 63% BiP cells) and (**D**) GFP^hi^ thymocytes (70% T, 30% BiP) during involution. *p*-values from 2-way ANOVA multiple comparison tests.

After preparing thymic single-cell suspensions, we analyzed cells within the lymphocyte gate (44) for GFP fluorescence intensity (Figure 2A). Strikingly, total GFP^+^ thymocytes in *lck*:*eGFP* fish declined ∼92% from ∼1.04 x 10^5^ thymocytes (or, since *D. rerio* have two thymi, ∼52K/thymus) at 3m to ∼8,500 GFP^+^ cells by 6m (*p*=0.0004; Figure 2B). By 12m, they declined >3-fold further, with >97% fewer thymocytes than prior to involution (∼2,700 thymocytes; *p*=0.0006) (Figure 2B and Table 1). The ratio of thymic GFP^hi^:GFP^lo^ cells also changed as involution progressed, with GFP^hi^ cells falling more precipitously than GFP^lo^ cells (Figure 2B, right histogram).

**Table 1:**
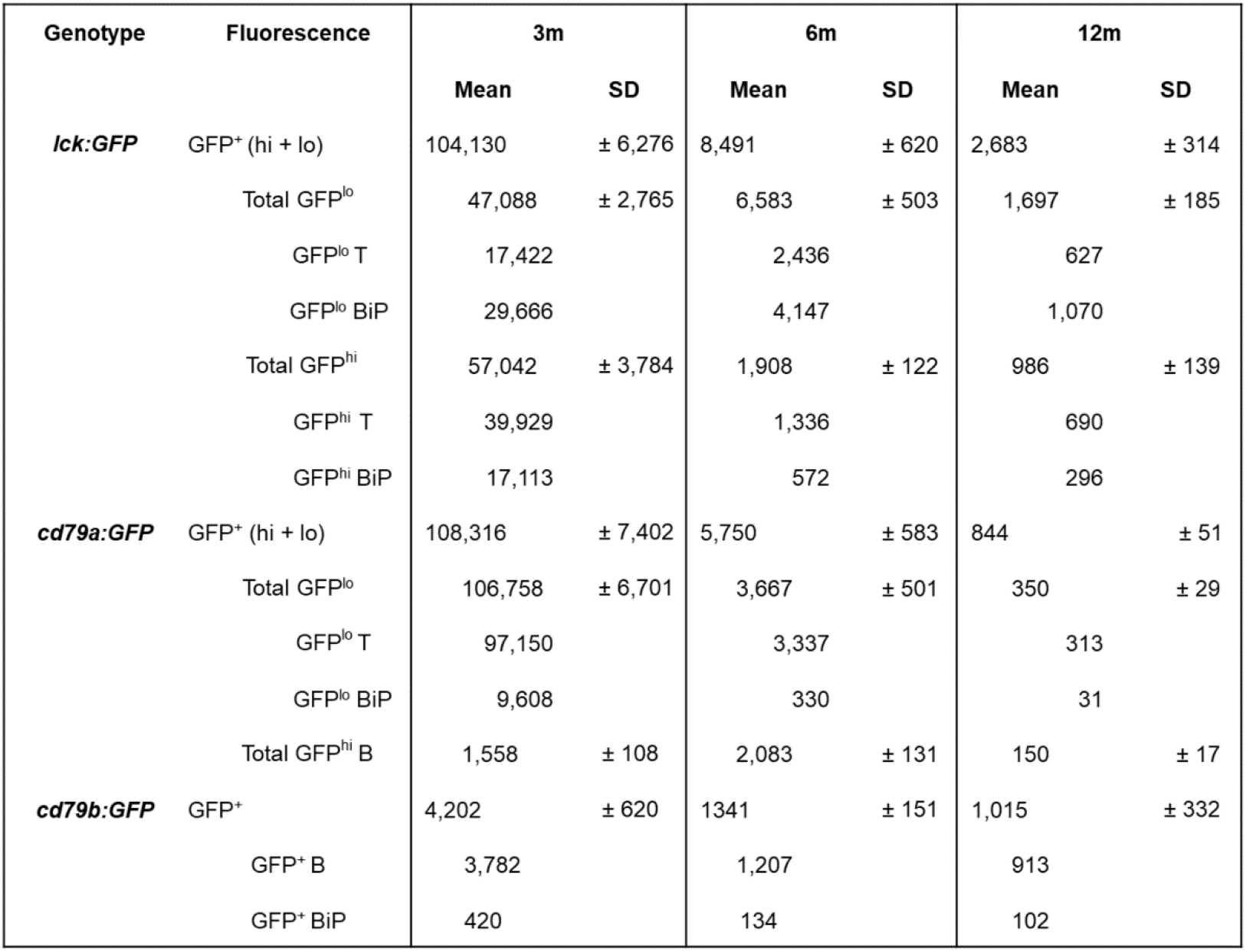
Thymic Lymphocyte Quantification.

| Genotype | Fluorescence | 3m |  | 6m |  | 12m |  |
| --- | --- | --- | --- | --- | --- | --- | --- |
|  |  | Mean | SD | Mean | SD | Mean | SD |
| ***lck:GFP*** | GFP<sup>+</sup> (hi + lo) | 104,130 | ± 6,276 | 8,491 | ± 620 | 2,683 | ± 314 |
|  | Total GFP<sup>lo</sup> | 47,088 | ± 2,765 | 6,583 | ± 503 | 1,697 | ± 185 |
|  | GFP<sup>lo</sup> T | 17,422 |  | 2,436 |  | 627 |  |
|  | GFP<sup>lo</sup> BiP | 29,666 |  | 4,147 |  | 1,070 |  |
|  | Total GFP<sup>hi</sup> | 57,042 | ± 3,784 | 1,908 | ± 122 | 986 | ± 139 |
|  | GFP<sup>hi</sup> T | 39,929 |  | 1,336 |  | 690 |  |
|  | GFP<sup>hi</sup> BiP | 17,113 |  | 572 |  | 296 |  |
| ***cd79a:GFP*** | GFP<sup>+</sup> (hi + lo) | 108,316 | ± 7,402 | 5,750 | ± 583 | 844 | ± 51 |
|  | Total GFP<sup>lo</sup> | 106,758 | ± 6,701 | 3,667 | ± 501 | 350 | ± 29 |
|  | GFP<sup>lo</sup> T | 97,150 |  | 3,337 |  | 313 |  |
|  | GFP<sup>lo</sup> BiP | 9,608 |  | 330 |  | 31 |  |
|  | Total GFP<sup>hi</sup> B | 1,558 | ± 108 | 2,083 | ± 131 | 150 | ± 17 |
| ***cd79b:GFP*** | GFP<sup>+</sup> | 4,202 | ± 620 | 1341 | ± 151 | 1,015 | ± 332 |
|  | GFP<sup>+</sup> B | 3,782 |  | 1,207 |  | 913 |  |
|  | GFP<sup>+</sup> BiP | 420 |  | 134 |  | 102 |  |

In *lck:eGFP* fish, GFP^+^ thymocytes show a wide range of GFP intensity spanning nearly two orders of magnitude (Figure 2A, right panel). Discrete GFP^+^ populations were not evident by flow cytometry, but gene expression profiles (GEP) vary across this GFP intensity spectrum. For example, we showed T- vs. B-ALL in *lck*:*eGFP* fish are GFP^hi^ vs. GFP^lo^, respectively (29). Our prior work also analyzed GFP^lo^ and GFP^hi^ fractions by qRT-PCR, with GFP^lo^ thymocytes showing B-lineage GEP and GFP^hi^ cells having T-lineage GEP (28); thus, we originally classified these as thymic B and T cells. Our recent single-cell qRT-PCR (sc-qRT-PCR; Park *et al.*, submitted; see attached pdf) data further refine this interpretation, with many GFP^lo^ thymocytes co-expressing B- and T-lineage genes. We refer to these cells as Bi-Phenotypic (BiP) lymphocytes. In contrast, far fewer GFP^hi^ thymocytes have BiP GEP. Using these data, we extrapolated cell type frequencies in the GFP^lo^ and GFP^hi^ fractions (Figure 2C-D).

GFP^lo^ thymocytes (comprising 37% T and 63% BiP cells) showed a similar pattern of decline from ∼4.7 x 10^4^ cells at 3m to ∼6,600 by 6m (86% fewer), with only ∼1,700 cells (>96% fewer) by 12m (Figure 2C and Table 1). Extrapolated declines in T and BiP cells were ∼17,400 → 2,400 → 600 and ∼30,000 → 4,100 → 1,000, respectively. GFP^hi^ thymocytes (70% T, 30% BiP) showed even higher rates of decline, from ∼5.7 x 10^4^ cells at 3m to ∼1,900 (>96% reduced) and ∼1,000 (>98% reduced) cells by 6 and 12m, respectively (Figure 2D and Table 1). These extrapolate to declines of ∼40,000 → 1,300 → 700 T cells and ∼17,000 → 600 → 300 BiP cells in the GFP^hi^ fraction.

The *cd79a:GFP* line was previously reported to be B-lineage specific (30). However, we found this to be inaccurate. Like *lck*:*eGFP* fish, *cd79a:GFP* thymocytes show varying GFP intensities, as well as distinct GFP^lo^ and GFP^hi^ populations (Figure 3A, right panel). These fractions also contain different percentages of T, B, and BiP cells by sc-qRT-PCR (Park *et al.*, submitted; see attached pdf). To quantify involution-related changes in *cd79a:GFP* thymocytes, we enumerated lymphoid gate GFP^lo^ and GFP^hi^ cells (Figure 3B-D). Total GFP^+^ thymocytes declined by >94%, from ∼10.8 x 10^4^ thymocytes at 3m to 5,800 by 6m (*p*=0.001); a further ∼7-fold decrease was seen by 12m (∼800 cells; *p*=0.001; Figure 3B and Table 1). Overall, by 12m, GFP^+^ thymocytes fell >98% from pre-involution, mirroring findings in *lck:eGFP* fish (Figure 2).

**Figure 3:**
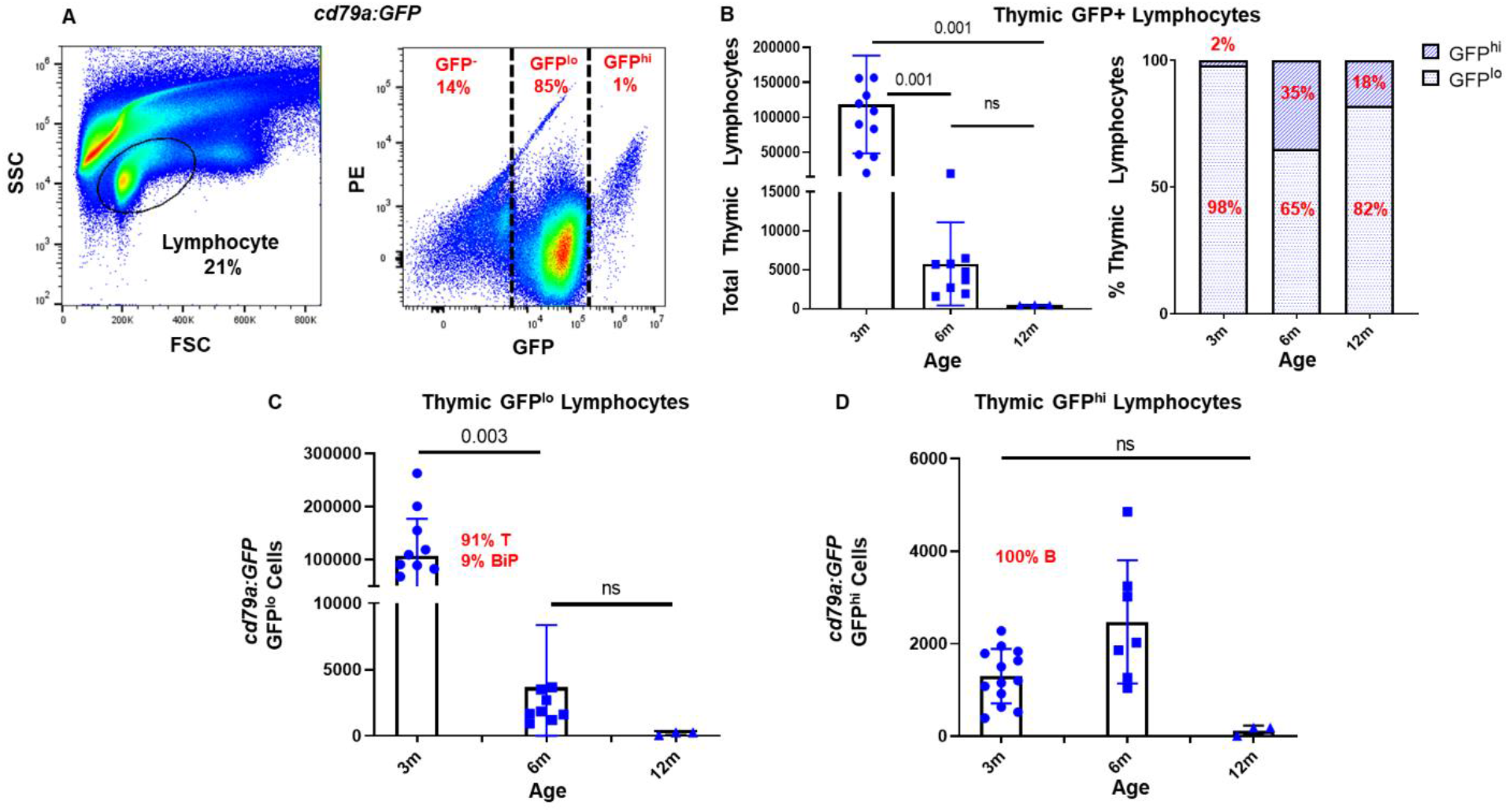
Quantification of thymocytes in *cd79a:GFP* zebrafish during involution. (**A**) Flow cytometry plots as in Figure 2. Note discrete GFP^lo^ and GFP^hi^ populations. (**B**) Quantification of total thymocytes (**left**) and % of GFP^lo^ vs. GFP^hi^ populations (**right**) at 3m (n=10 fish), 6m (n=9), and 12m (n=3). Declining (**C**) GFP^lo^ (91% T, 9% BiP cells) and (**D**) statistically-stable GFP^hi^ thymocytes (100% B) during involution. *p*-values from 2-way ANOVA multiple comparison tests.

GFP^lo^ cells in *cd79a:GFP* fish declined significantly, from ∼10.7 x 10^4^ cells at 3m to ∼3,700 by 6m (>96% less) and ∼350 cells by 12m (>99% less; Figure 3C and Table 1). GFP^lo^ thymocytes are comprised of 91% T and 9% BiP cells (Park *et al.*, submitted; see attached pdf). Extrapolating from these percentages, GFP^lo^ thymocytes had estimated declines of 97,000 → 3,300 → 300 and 9600 → 300 → 30 for T and BiP cells, respectively (Figure 3C, Table 1). GFP^hi^ thymocytes are purely B-lineage in *cd79a:GFP* (Park *et al.*, submitted; see attached pdf). Intriguingly, thymic B cells rose slightly during involution, from ∼1,600 to ∼2,100 cells during the 3-6m window. However, by 12m, thymic B cells declined to 150 cells (90% reduced; Figure 3D and Table 1). Thymic B cell absolute numbers did fall post-involution, but since T and BiP declines were more profound, thymic B cells actually increased relative to these. In fact, by 12m, 18% of total thymocytes were B-lineage (Figure 3B, right panel). Humans also have a relative increase in thymic B cells post-involution (13), suggesting this phenomenon is conserved across vertebrates. We also analyzed data based on sex, which showed no differences between female and male involution kinetics in either the *lck*:*eGFP* or *cd79a*:*GFP* lines (Supplemental Figure 1).

*cd79b:GFP* fish have only one thymic GFP^+^ population (Figure 4A, right panel), consisting of 90% B and 10% BiP cells (Park *et al.*, submitted; see attached pdf). GFP^+^ cells declined from ∼4,200 cells at 3m to ∼1,300 cells (68% less) and ∼1,000 (76% less) by 6 and 12m, respectively (Figure 4B, Table 1). These extrapolate to declines of ∼3,800 → 1,200 → 900 B cells and 420 → 130 → 100 BiP cells. Notably, we detected more thymic B cells in *cd79b:GFP* than *cd79a:GFP* fish, suggesting the former labels B-lineage cells more completely. In summary, flow cytometric studies of *lck:eGFP*, *cd79a:GFP*, and *cd79b:GFP* thymocytes reveal consistent and marked decline in thymic lymphocytes during the 3-6m window, congruent with decreasing thymic fluorescence caused by involution in that same time period.

**Figure 4:**
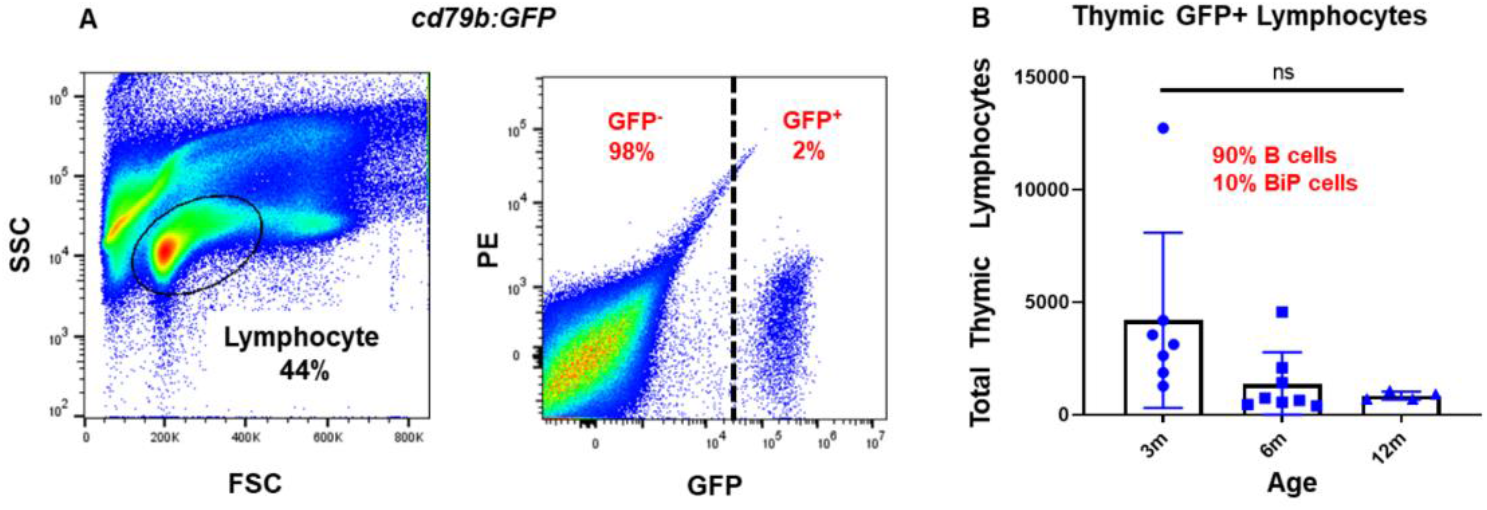
Quantification of thymocytes in *cd79b:GFP* zebrafish during involution. (**A**) Flow cytometry plots as in Figure 2. Note single GFP^+^ population. (**B**) Quantification of total GFP^+^ thymocytes (90% B, 10% BiP cells) at 3 (n=7 fish), 6 (n=8), and 12m (n=4).

### Changes in Thymic Morphology During Involution

Having observed diminishing fluorescent signals by microscopy and declining thymocytes by flow cytometry, we next analyzed thymic morphology in WT fish. To do this, we serially sectioned fish, performed H&E stains, measured thymic areas, and calculated thymic volumes for 3, 6, and 12m WT fish. On sagittal view, the thymus is caudal and dorsal to the eye (Figure 5A, left panel). We performed serial sagittal sectioning beginning at each animal’s surface, proceeding lateral → medial, as diagrammed on coronal views (Figure 5A, center and right panels), with H&E staining at 20µm intervals. Three-month zebrafish thymi had well-defined cortical and medullary regions (Figure 5B, top right panel, yellow dashed line). By 6m, thymi were markedly smaller with less distinct cortico- medullary junctions (Figure 5B, middle row). By 12m, cortex and medulla were indistinct in thymic remnants (Figure 5B, bottom row). This coincided with adipose replacement of thymic tissue, which also occurs in mammals (45) (black arrow in Figure 5B lower right panel).

**Figure 5:**
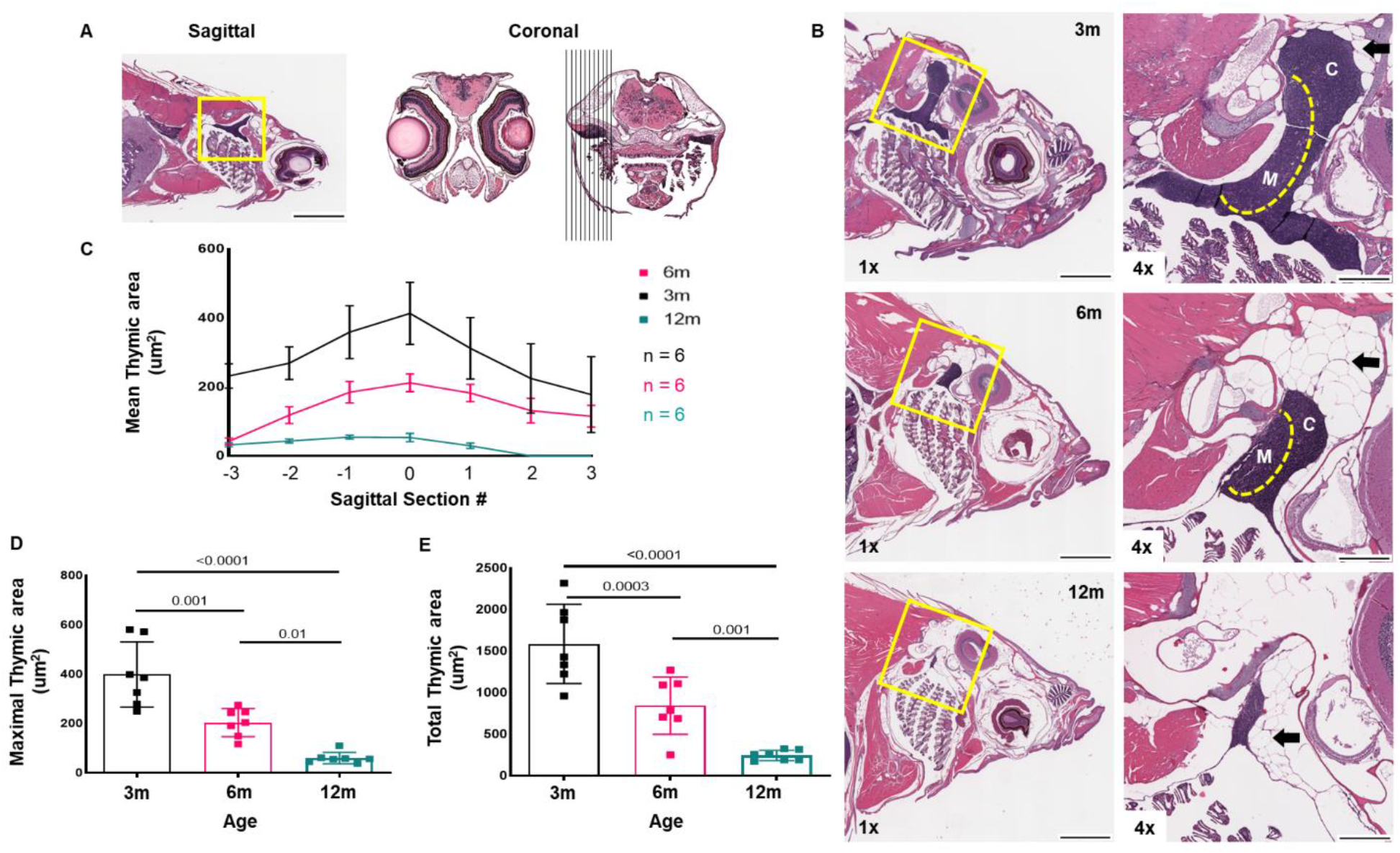
Morphologic changes in zebrafish thymi with involution. (**A**) Sagittal section; yellow box highlights thymus (**L**). Schematic of serial coronal sections to determine thymic areas (**R**). (**B**) H&E stains of 3 (top), 6 (middle), and 12m (bottom) zebrafish demonstrating progressive involution. (**L**) 1X (scale bar = 1mm), (**R**) 4X (scale bar = 0.25 mm); labels indicate cortex (**C**), medulla (**M**), and peri-thymic adipose tissue (**black arrows**). (**C**) Mean thymic areas (hashes denote SEM) across serial sections, at 3, 6, and 12m. (**D**) Maximal and (**E**) Total thymic areas at 3, 6, and 12m. For each group, n = 6, units in µm^2^; *p*-values by 2-way ANOVA multiple comparison tests.

To quantify changes in thymic size, we calculated mean and maximal thymic areas for each age by defining thymic boundaries on H&E-stained sections via ImageJ (Figure 5C-D). To calculate mean areas, we used 7 consecutive slides from each fish (total depth = 120µm) that contained the largest amount of thymic tissue. This captured the entire thymus in 6 and 12m fish, and the bulk of the thymus in 3m fish (Figure 5C). At every point of comparison, 3m thymic areas exceeded 6m fish by ∼2-fold; similarly, 6m thymic areas were much larger than 12m thymi (Figure 5C). Maximal and total (summed across the 120µm span) thymic areas at 3, 6, and 12m are depicted in Figures 5D-E, respectively. Comparing maximal (Sagittal Section “0” in Figure 5C) and total thymic areas yielded similar estimates of involution-induced changes, with 3m thymi having mean maximal thymic areas of 400 µm^2^, which decreased by 50% to 200 um^2^ by 6m (*p*<0.001). By 12m, mean maximal thymic area was ∼3.3-fold smaller at 60 µm^2^ (85% reduced from 3m; *p*=0.0001; Figure 3D). We extrapolated thymic volumes to build 3D models of representative thymi at each age (Figure 6, [20230614.3m.6m.12m.thymus.models.avi](https://sooners-my.sharepoint.com/:v:/g/personal/clay-foster_ouhsc_edu/EbKj-1GTKthKr0AN-RTcgFIBPrOpNraKO-Po_m9IB2cbJQ?e=sZaJ5V)).

**Figure 6:**
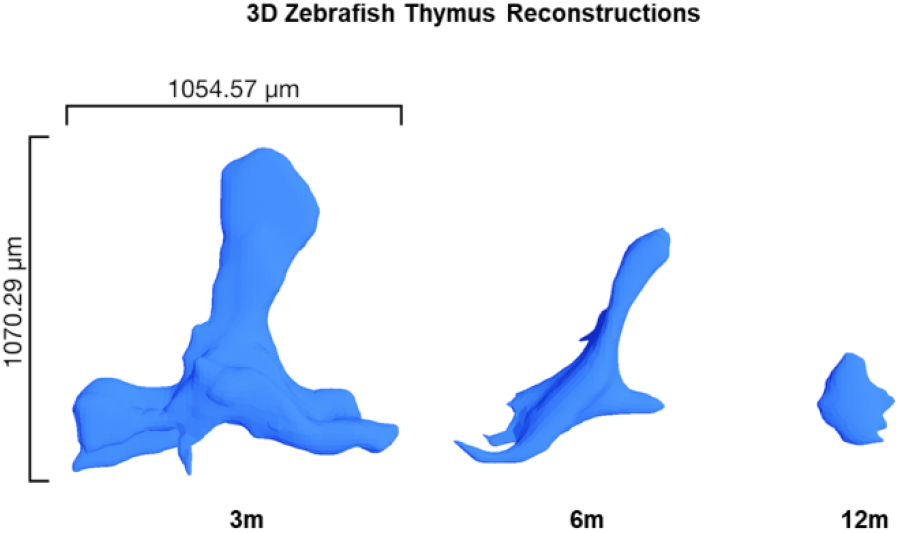
3D zebrafish thymi reconstructions at different involution timepoints. 3D reconstructions of zebrafish thymi at 3, 6, and 12m. Volumetric estimates of each thymus are: 3.5 x 10^6^ µm^3^, 1.7 x 10^6^ µm^3^, and 3.2 x 10^5^ µm^3^, respectively.

Absolute thymus volumes were 3.52 x 10^6^ µm^3^ at 3m, 1.70 x 10^6^ µm^3^ (52% smaller) at 6m, and 3.32 x 10^5^ µm^3^ (91% decreased from original) at 12m. To normalize for growth, we also calculated thymus:brain volumetric ratios, which revealed a pattern of age-related atrophy (0.33 → 0.11 → 0.013) over the 9m span. These morphologic data, together with fluorescent microscopy and flow cytometry results (Figures 1-4), show structural and cellular changes of thymic involution that support the proposed Figure 1B timeline, with marked involution during 3-6m and continued regression over the following 6 months.

### Expression Signatures of Thymic and Marrow Lymphocyte Subsets

We next sought to identify GEP for the different lymphocyte populations impacted by involution. To do this, we built four novel double-transgenic lines to fractionate T and B cells into highly-refined subsets for bulk RNA sequencing. TCR and immunoglobulin (Ig) rearrangement is mediated by recombination-activating gene *rag1* and *rag2* protein products (46, 47). Thus, *rag1*/*2* expression is a proxy for immature lymphoblasts, making *rag2:RFP* transgenic fish (where a *D. rerio rag2* promoter regulates RFP) a lymphoblast- specific marker line. We bred this line to *cd79a:GFP* and *cd79b:GFP* fish to build double-transgenic fish with both stage- (immature vs. mature lymphocyte) and lineage- (B vs. T) specific markers. Liu *et al.* previously employed a similar dual-transgene approach (*rag2:mCherry* plus *cd79a*:*GFP* or *cd79b*:*GFP*), but only analyzed marrow B cells, and only tested seven transcripts by qRT-PCR (30). We also bred *lck:mCherry* to *cd79a:GFP* and *cd79b:GFP* fish to make two novel lines with distinct T and B cell fluorescent labels. Using these markers, we FACS-purified multiple thymic (Figure 7, 1^st^ column) and marrow (Figure 7, 2^nd^ column) populations from each double-transgenic. We performed RNA-seq of each in triplicate, seeking differentially-expressed genes in each lymphocyte subset, including thymic B cells (T4). Principal Component Analysis (PCA) of thymic and marrow lymphocyte subsets from *rag2*:*RFP*;*cd79a*:*GFP* (Figure 7A, right panel) and *rag2*:*RFP*;*cd79b*:*GFP* (Figure 7B, right panel) resolved distinct clusters, with T-lineage subsets at left and B-subsets (including thymic B) at right (Figure 7C depicts a merged PCA for both genotypes). Likewise, PCA of subsets from *lck*:*mCherry*;*cd79a*:*GFP* (Figure 7D, right panel) and *lck*:*mCherry*;*cd79b*:*GFP* (Figure 7E, right panel) clustered T (T1, T2), B (T4), and immature (M1) lymphocytes (merged PCA for both genotypes in Figure 7F; merged PCA for all four genotypes in Supplemental Figure 2A). Notably, thymic B cells (T4) of all four genotypes were near-superimposable by PCA, and aligned closely to marrow B subsets (Figure 7C, 7F and Supplemental Figure 2A).

**Figure 7:**
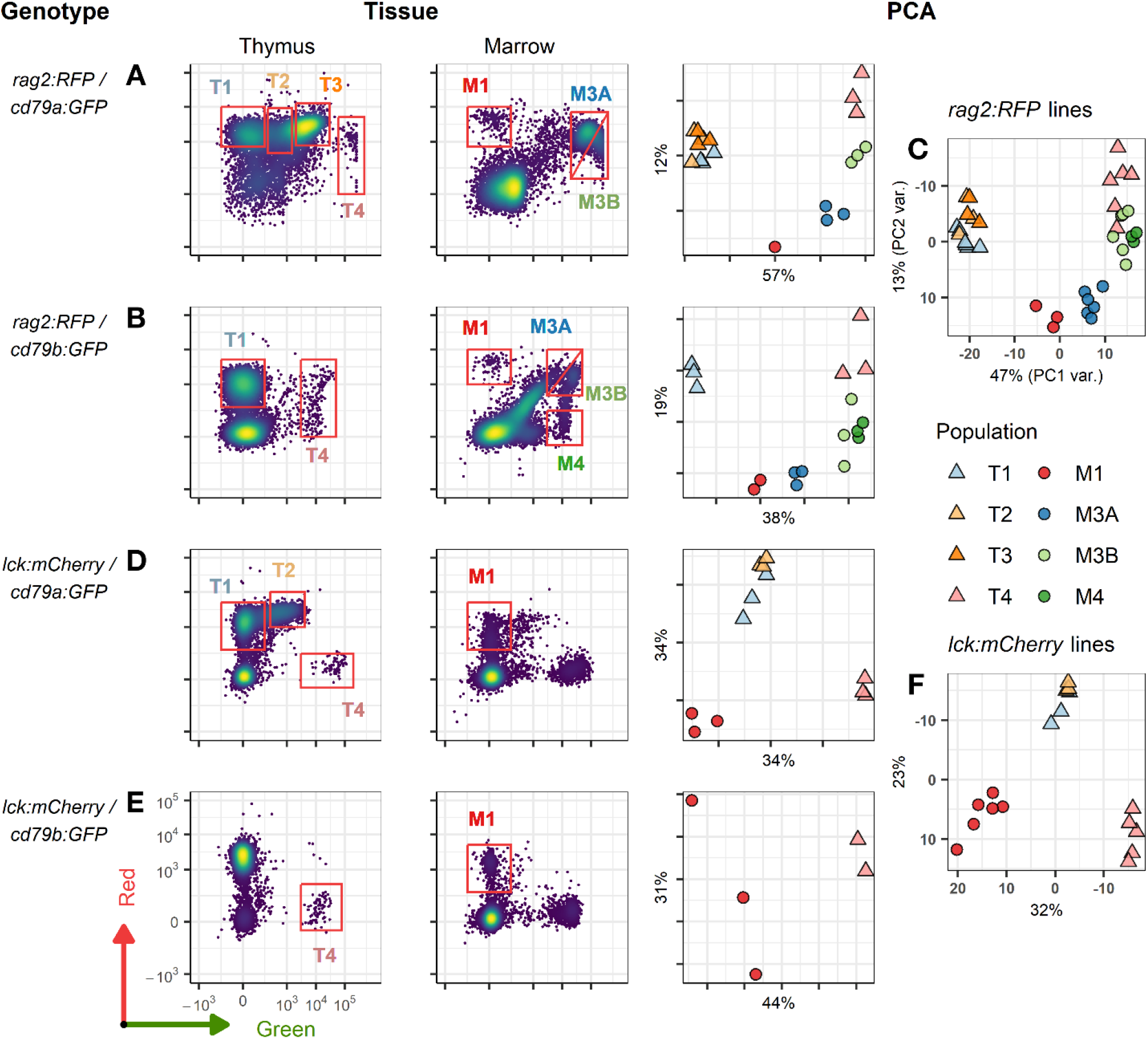
FACS profiles of thymic and marrow lymphocyte subsets and PCA cluster analyses. (**A**) FACS plots of double-transgenic, *rag2*:*RFP* + *cd79a*:*GFP*, thymic (left column) and marrow (middle column) lymphocytes. Red boxes show gates of populations analyzed by bulk RNA-seq as triplicate (T1-T4, M3A, M3B) or single (M1) samples. Principal Component Analysis (PCA using top 1000 most variable genes; right column) shows clustering of sample types. (**B**) Same depiction as in **A** for *rag2*:*RFP* + *cd79b*:*GFP* thymic and marrow lymphocytes of triplicate- (T1, T4, M3A, M3B, M4) and duplicate- (M1) sequenced samples. (**C**) PCA of all 36 samples from both *rag2*:*RFP* lines in **A** and **B**. (**D**) Same depiction as above for double-transgenic, *lck*:*mCherry* + *cd79a*:*GFP*, thymic and marrow lymphocytes sequenced in triplicate (T1, T2, T4, M1). (**E**) Same depiction as above for *lck*:*mCherry* + *cd79b*:*GFP* thymic and marrow lymphocytes sequenced in duplicate (T4) or triplicate (M1). (**F**) PCA of all 17 samples from both *lck*:*mCherry* lines in **D** and **E**.

Unbiased analysis of subsets from all four double-transgenic lines using the 1000 most-variable genes revealed distinct GEP that clearly distinguished T- vs. B-lineage cells (Figure 8A; Supplemental Table 1 lists all genes in this heatmap, which positively and negatively correlate with PC1 to separate T vs. B). To more closely examine subsets of each lineage, which T vs. B differences mask when comparing all samples together, we next analyzed T- (Figure 8A, left and 8B PCA) and B- (Figure 8A, right and 8C PCA) subsets separately, seeking distinct maturational stages (3^rd^ row of Figure 8B-C and Supplemental Figure 2B-C). This strategy excludes pan-lineage markers like *cd4*, *cd8*, *cd79a*/*b*, etc. to find differentially-expressed genes within a specific lineage.

**Figure 8:**
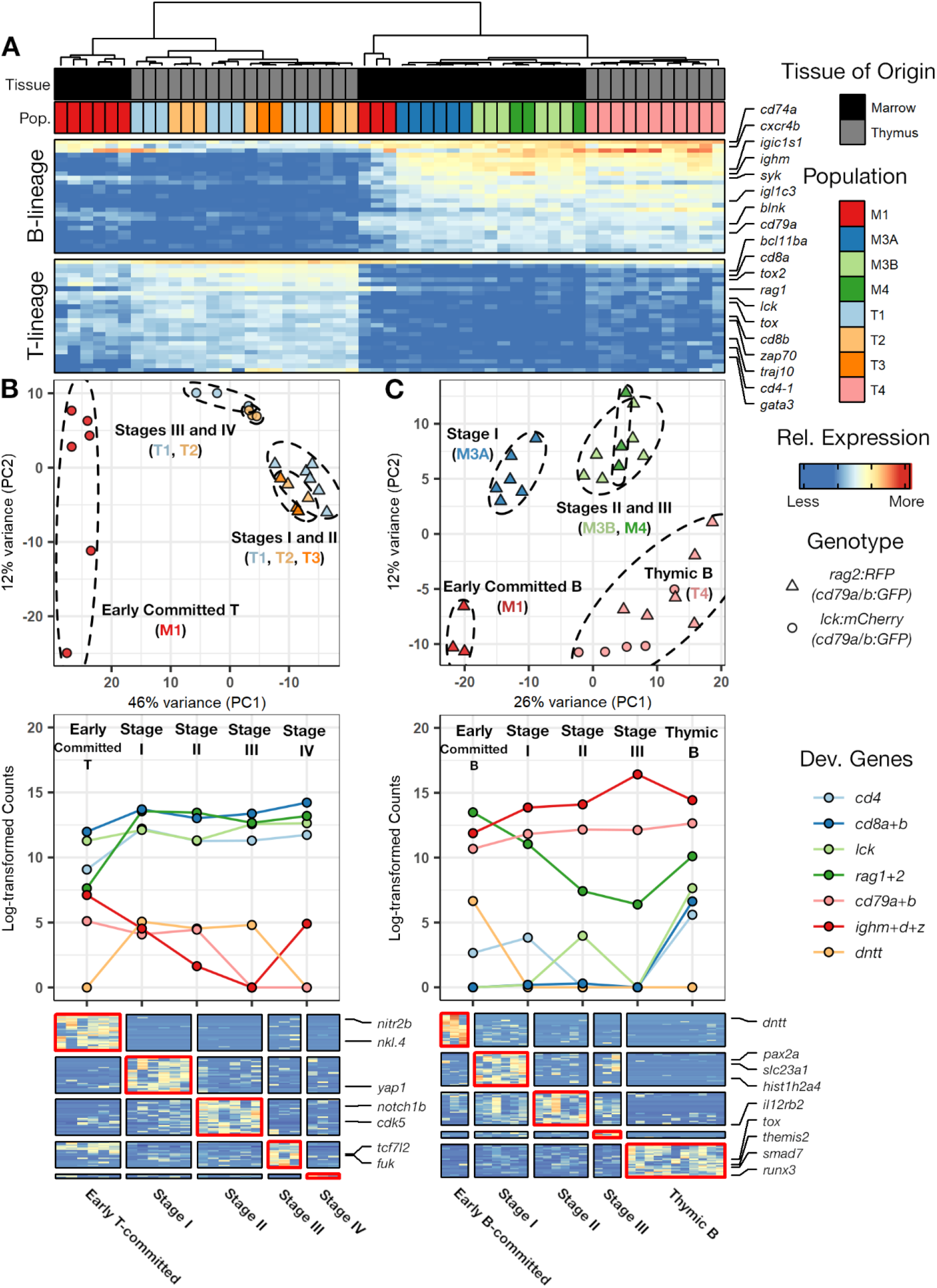
Expression profiles of thymic and marrow lymphocyte subsets. (**A**) Expression of the top 50 genes distinguishing T- vs. B-lineage thymic and marrow lymphocytes (correlated with PC1 as seen in Figure S3A). Annotations at top correspond to tissue-of-origin (black = marrow, grey = thymus) and color-coding of populations collected by FACS. (**B**) Top: PCA of T-lineage subsets clusters distinct T maturational stages; Total samples (n=24), Early-committed T (*lck:mCherry* M1, red circles in PCA; n=6), Stage I (*rag2*:*RFP*;*cd79a*:*GFP* T2 and T3, light- and dark-orange triangles in PCA; n=6), Stage II (*rag2*:*RFP*;*cd79a* or *cd79b*:*GFP* T1, light-blue triangles in PCA; n=6), Stage III (*lck:mCherry*;*cd79a*:*GFP* T1, light-blue circles in PCA; n=3), and Stage IV (*lck:mCherry*;*cd79a*:*GFP* T2, light-orange circles in PCA; n=3). Middle: normalized counts of T-lymphopoiesis genes in T-lineage subsets. Bottom: subset-specific genes for each T-lymphopoietic stage. (**C**) Identical depictions for B-lineage samples, including Thymic B cells. Total samples (n=29), Early-committed B (*rag2*:*RFP* M1, red triangles in PCA; n=3), Stage I (*rag2*:*RFP*;*cd79a*:*GFP* or *cd79a*:*GFP* M3A, blue triangles in PCA; n=6), Stage II (*rag2*:*RFP*;*cd79a*:*GFP* or *cd79a*:*GFP* M3B, light-green triangles in PCA; n=6), Stage III (*rag2*:*RFP*;*cd79b*:*GFP* M4, dark-green triangles in PCA; n=3), and thymic B cells (all four genotypes’ T4, pink triangles and circles in PCA; n=11).

T-lineage subsets (M1 from *lck*:*mCherry*, T1, T2, T3) expressed high levels of *cd4*, *cd8a/b, lck, rag1*/*2*, with the only marrow T-subset (M1) showing slightly lower levels (Figure 8B, 3^rd^ row). M1 also had higher Ig (*ighm*, *ighd*, *ighz*/*igt*) and *cd79a*/*b* transcripts, suggesting this immature T-lineage cluster is still extinguishing B-lineage expression as these cells prepare to transit from marrow to thymus. Other genes linked to mammalian T-lymphopoiesis [*cd34*, *gata3*, *dntt* (i.e., TdT), *her6* (homologous to *HES1*), etc.] also exhibited expression trajectories supporting our proposed T cell differentiation schema (Supplemental Figure 2B). Genes unique-expressed by each T cell developmental stage are shown in the bottom panel of Figure 8B (complete genelist, Supplemental Table 1).

B-lineage subsets (M1 from *rag2*:*RFP*, M3A, M3B, M4, and T4) expressed high Ig and *cd79* transcripts, with M1 cells having highest *rag1*/*2* and *dntt*, implying M1 are the earliest B-precursors (Figure 8C, 3^rd^ row). Conversely, subsequent B cell stages had progressively lower *rag1*/*2* and *dntt*, while their Ig and *cd79a*/*b* progressively increased. Other genes linked to mammalian B-lymphopoiesis [*cd34*, *sox4a*, *syk*, *ptprc* (i.e., cd45), etc.] likewise displayed expression patterns supporting our B cell differentiation schema (Supplemental Figure 2C). Genes unique to each B cell developmental stage are shown in the bottom panel of Figure 8C (complete genelist, Supplemental Table 1).

We also compared thymic B cells (T4) of all four double-transgenic lines to marrow stage III B cells (M4). These groups cluster closely by PCA (Supplemental Figure 2A), but we identified 312 genes whose expression differentiates thymic vs. marrow B cells (Supplemental Figure 2D, Supplemental Table 1). This represents the first description of thymic B cell-specific markers in zebrafish. One notable thymic B cell feature was their expression of “T cell genes” (*rag1*, *tox*, *tox2*) that also differentiated the T- vs. B-lineages (Figure 8A). This could suggest contamination of T4 by T cells, although T4 cells were far removed from other subsets in FACS collections (left plots of Figure 7A-B, D-E). To exclude this possibility, we examined the 25 transcripts uniquely up-regulated by thymic B cells vs. all other B-subsets (Figure 8C, bottom right heatmap), excluding any that were highly expressed by any T-subset (M1 from *lck*:*mCherry*, T1, T2, T3). This revealed 13 genes that distinguish thymic B cells from *every other lymphocyte subset* (Supplemental Figure 2E). This stringent requirement assures these are *bona fide* thymic B markers, although many excluded genes are unlikely due to T contamination, which we will test by single-cell methods going forward.

We also examined Ig transcripts in thymic B cells, which revealed they express far more *ighz* (alternatively designated IgT) than marrow B cells (Supplemental Figure 3; 11 thymic B vs. 3 stage III marrow B samples, *p*-value = 0.058). IgZ/IgT is an Ig class for mucosal immunity, functionally like mammalian IgA (48). However, unlike in mammals, *ighz* expression does not occur by Ig class-switching, rather, it is produced by a distinct B cell lineage (49). Thus, our results imply the thymus may be the site of IgZ-B lymphopoiesis, or possibly the primary site of IgZ-B cells. Conversely, stage III marrow B cells expressed more *ighm* than thymic B cells (*p*-value = 0.12), with no noteworthy differences in *ighd* or Ig-light chain transcripts. Overall, these thymic B cell transcriptomic profiles contribute to our understanding of *D. rerio* B-lymphopoiesis, enhancing zebrafish as a vertebrate adaptive immunity model.

## Discussion

Over the past ∼30 years, zebrafish have gradually emerged as a model to study vertebrate lymphopoiesis and immunity. Extensive work in *D. rerio* provides compelling evidence that thymopoiesis and T cell development are evolutionarily conserved from teleost fish to mammals (4, 50). Most zebrafish studies have utilized embryonic and larval stage fish, largely due to imaging advantages, since these early stages are transparent-to-translucent. However, there is growing recognition of the importance of immune studies in adult zebrafish. For example, since thymic involution occurs later in life, this phenomenon cannot be interrogated in immature zebrafish.

The thymus is not static; in humans, it changes throughout the entire lifespan (51). Thymic involution is just one component of immunosenescence, but likely the earliest, since the majority of human involution occurs during adolescence (45). If human and *D. rerio* involution are similar, zebrafish provide a useful tool for involution studies, because they are genetically-tractable in terms of their amenability to transgenesis, CRISPR/Cas9, and related manipulations. However, studies of zebrafish thymic involution were virtually non-existent, with the original description of the *lck*:*eGFP* line we used here noting that thymic fluorescence diminished with aging (21), and a more recent study examining thymic structural/morphometric changes in WT and *lck*:*GFP* fish (20). This later work concluded that zebrafish thymic involution coincides with attainment of sexual maturity, like humans. However, it did not address changes in thymocyte numbers during involution, focusing only the organ as a whole.

In the current work, we expanded upon this, testing multiple lymphocyte-labeled genotypes (Figure 1), using new morphologic quantification strategies (Figures 5-6), and critically, enumerating the specific numbers of different thymic lymphocytes pre- and post-involution (Figures 2-4). By every metric, the majority of thymic atrophy occurred in the 3- 6m window, and by 12m, <10% of thymic fluorescence, thymocyte numbers, and thymic area/volume remained. In aggregate, these results support the hypothetical involution timeline we propose in Figure 1B. We also demonstrated that thymic B cells, a curious population whose role(s) are not fully known, are less susceptible to involution-mediated decline than thymic T cells, although involution still reduces them markedly, as in humans. This and other conserved features between zebrafish and human involution not only yield insight into the biology underlying thymic immunosenescence, but also highlight the potential of zebrafish models to enable studies on the pathways and mechanisms regulating thymic involution and how it impacts immune function overall.

We also made and analyzed four novel double-transgenic zebrafish models where lymphoblast, B-, and/or T-lineage cells are differentially labeled. These lines are valuable to the field, expanding the specificity and types of experiments that can be done in zebrafish adaptive immunity studies. We used RNA-seq to identify several distinct T- and B- maturational stages in both lymphopoietic organs, marrow and thymus, of these lines. Previously, B cell maturation in zebrafish marrow was examined using dual-transgenic *rag2*:*mCherry*;*cd79*:*GFP* fish, but this report did not evaluate thymocytes, and was limited to bulk qRT-PCR of seven transcripts (*ighm*, *ighz*, *igic1s1*, *cd79a*, *cd79b*, *rag1*, *rag2*) (30). Our inclusion of thymic lymphocytes and complete transcriptomic profiling by RNA-seq expands upon their findings, largely supporting their B-lymphopoiesis schema with additional refinements. We also identified lineage- and stage-specific transcripts for every T and B cell maturation stage (Figure 8B, bottom panels).

Knowledge about zebrafish thymic B cells is limited. Their existence was first shown by Liu *et al*. using fluorescent microscopy to visualize B cells on the surface of the thymus (30). Intriguingly, our prior work also found that B-ALL often envelop the thymic surface (29). A recent study reported a ‘transcriptional atlas’ for zebrafish marrow and thymus compiled by single-cell RNA-seq (scRNA-seq) (22). They analyzed 530 thymic and 3,656 marrow B cells, identifying distinct zebrafish B cell developmental stages. Here, we also characterized maturation of marrow and thymic B cells and T cells. While our approach lacks single-cell resolution, our use of bulk RNA-seq to analyze multiple fluorescently-distinct populations gives much deeper transcriptomic data than scRNA-seq can achieve. Moreover, by sequencing RNA from ∼200,000 thymic B cells (11 independent replicates from 4 genotypes), we obtained a comprehensive profile that complements and expands upon the initial 530 thymic B cells derived from scRNA-seq.

We identified 312 genes that distinguish thymic versus marrow B cells, providing a starting point to investigate their unique functions. One key difference may be higher *ighz* expression by thymic B cells. Supporting this, thymic B-ALL in our zebrafish model were consistently IgZ-lineage (52). IgZ is important for mucosal immunity, and the thymus is anatomically near both the gills and oropharynx—sites where pathogens contact the mucosae. Thus, zebrafish thymic B cells may have specialized function in this regard. Overall, this work highlights the diversity of lymphocyte populations in zebrafish thymus, including thymic B cells. In addition, by investigating thymic involution, we establish zebrafish as a potentially-powerful involution model. Future investigations of thymic B cell function and the mechanisms governing thymic involution can enhance our understanding of these evolutionarily-conserved phenomena in humans.

## Supporting information

STable 1

## Acknowledgments

We thank Louisa Williams and Jim Henthorn in the OUHSC tissue histology and flow cytometry cores, respectively, for their expertise and contributions to our experiments. We thank the Langenau lab (Massachusetts General Hospital, Harvard Medical School, and Harvard Stem Cell Institute) for *rag2*:*RFP* fish, the Zon lab (Boston Children’s Hospital, Dana-Farber Cancer Institute, Harvard University, HHMI, Harvard Medical School, and Harvard Stem Cell Institute) for lck:mCherry fish, and the Hardy lab (Fox Chase Cancer Center, Temple Health) for cd79a:GFP and cd79b:GFP fish.

## References

1. Boehm, T., and J. B. Swann. 2013. Thymus involution and regeneration: Two sides of the same coin? Nat Rev Immunol 13: 831–838.

2. Hale, L. P. 2004. Histologic and molecular assessment of human thymus. Ann Diagn Pathol 8: 50–60.

3. Shanley, D. P., D. Aw, N. R. Manley, and D. B. Palmer. 2009. An evolutionary perspective on the mechanisms of immunosenescence. Trends Immunol 30: 374–381.

4. Bajoghli, B., A. M. Dick, A. Claasen, L. Doll, and N. Aghaallaei. 2019. Zebrafish and medaka: Two teleost models of T cell and thymic development. Int J Mol Sci 20.

5. Lynch, H. E., G. L. Goldberg, A. Chidgey, M. R. Van den Brink, R. Boyd, and G. D. Sempowski. 2009. Thymic involution and immune reconstitution. Trends Immunol 30: 366–373.

6. Sempowski, G. D., L. P. Hale, J. S. Sundy, J. M. Massey, R. A. Koup, D. C. Douek, D. D. Patel, and B. F. Haynes. 2000. Leukemia inhibitory factor, oncostatin M, IL-6, and stem cell factor mRNA expression in human thymus increases with age and is associated with thymic atrophy. J Immunol 164: 2180–2187.

7. George, A. J. T., and M. A. Ritter. 1996. Thymic involution with ageing: Obsolescence or good housekeeping? Immunol Today 17: 267–272.

8. Posnett, D. N., D. Yarilin, J. R. Valiando, F. Li, F. Y. Liew, M. E. Weksler, and P. Szabo. 2003. Oligoclonal expansions of antigen-specific CD8(+) T cells in aged mice. Ann NY Acad Sci 987: 274–279.

9. Goronzy, J. J., W. W. Lee, and C. M. Weyand. 2007. Aging and T cell diversity. Exp Gerontol 42: 400–406.

10. Linton, P. J., and K. Dorshkind. 2004. Age-related changes in lymphocyte development and function. Nat Immunol 5: 133–139.

11. Dixit, V. D. 2010. Thymic fatness and approaches to enhance thymopoietic fitness in aging. Curr Opin Immunol 22: 521–528.

12. Steinmann, G. G., B. Klaus, and H. K. Mullerhermelink. 1985. The involution of the aging human thymic epithelium is independent of puberty - A morphometric study. Scand J Immunol 22: 563–575.

13. Nunez, S., C. Moore, B. Gao, K. Rogers, Y. Hidalgo, P. J. Del Nido, S. Restaino, Y. Naka, G. Bhagat, J. C. Madsen, M. R. Bono, and E. Zorn. 2016. The human thymus perivascular space is a functional niche for viral-specific plasma cells. Sci Immunol 1.

14. Perera, J., and H. Huang. 2015. The development and function of thymic B cells. Cell Mol Life Sci 72: 2657–2663.

15. Le, J., J. E. Park, V. L. Ha, A. Luong, S. Branciamore, A. S. Rodin, G. Gogoshin, F. Li, Y. E. Loh, V. Camacho, S. B. Patel, R. S. Welner, and C. Parekh. 2020. Single-cell RNA-seq mapping of human thymopoiesis reveals lineage specification trajectories and a commitment spectrum in T cell development. Immunity 52: 1105–1118 e1109.

16. Yamano, T., J. Nedjic, M. Hinterberger, M. Steinert, S. Koser, S. Pinto, N. Gerdes, E. Lutgens, N. Ishimaru, M. Busslinger, B. Brors, B. Kyewski, and L. Klein. 2015. Thymic B cells are licensed to present self antigens for central T cell tolerance induction. Immunity 42: 1048–1061.

17. Cockburn, A. 1992. Evolutionary ecology of the immune system - Why does the thymus involute. Funct Ecol 6: 364–370.

18. Contreiras, E. C., H. L. Lenzi, M. N. Meirelles, L. F. Caputo, T. J. Calado, D. M. Villa-Verde, and W. Savino. 2004. The equine thymus microenvironment: A morphological and immunohistochemical analysis. Dev Comp Immunol 28: 251–264.

19. Peel, E., and K. Belov. 2017. Immune-endocrine interactions in marsupials and monotremes. Gen Comp Endocrinol 244: 178–185.

20. Kernen, L., J. Rieder, A. Duus, H. Holbech, H. Segner, and C. Bailey. 2020. Thymus development in the zebrafish (*Danio rerio*) from an ecoimmunology perspective. J Exp Zool A Ecol Integr Physiol 333: 805–819.

21. Langenau, D. M., A. A. Ferrando, D. Traver, J. L. Kutok, J. P. Hezel, J. P. Kanki, L. I. Zon, A. T. Look, and N. S. Trede. 2004. *In vivo* tracking of T cell development, ablation, and engraftment in transgenic zebrafish. Proc Natl Acad Sci USA 101: 7369–7374.

22. Rubin, S. A., C. S. Baron, C. Pessoa Rodrigues, M. Duran, A. F. Corbin, S. P. Yang, C. Trapnell, and L. I. Zon. 2022. Single-cell analyses reveal early thymic progenitors and pre-B cells in zebrafish. J Exp Med 219.

23. Paik, E. J., and L. I. Zon. 2010. Hematopoietic development in the zebrafish. Int J Dev Biol 54: 1127–1137.

24. Danilova, N., V. S. Hohman, F. Sacher, T. Ota, C. E. Willett, and L. A. Steiner. 2004. T cells and the thymus in developing zebrafish. Dev Comp Immunol 28: 755–767.

25. Dee, C. T., R. T. Nagaraju, E. I. Athanasiadis, C. Gray, L. Fernandez Del Ama, S. A. Johnston, C. J. Secombes, A. Cvejic, and A. F. Hurlstone. 2016. CD4-transgenic zebrafish reveal tissue-resident Th2- and regulatory T cell-like populations and diverse mononuclear phagocytes. J Immunol 197: 3520–3530.

26. Lam, S. H., H. L. Chua, Z. Gong, Z. Wen, T. J. Lam, and Y. M. Sin. 2002. Morphologic transformation of the thymus in developing zebrafish. Dev Dyn 225: 87–94.

27. Langenau, D. M., and L. I. Zon. 2005. The zebrafish: A new model of T cell and thymic development. Nat Rev Immunol 5: 307–317.

28. Burroughs-Garcia, J., A. Hasan, G. Park, C. Borga, and J. K. Frazer. 2019. Isolating malignant and non-malignant B Cells from *lck:eGFP* zebrafish. J Vis Exp.

29. Borga, C., G. Park, C. Foster, J. Burroughs-Garcia, M. Marchesin, R. Shah, A. Hasan, S. T. Ahmed, S. Bresolin, L. Batchelor, T. Scordino, R. R. Miles, G. te Kronnie, J. L. Regens, and J. K. Frazer. 2019. Simultaneous B and T cell acute lymphoblastic leukemias in zebrafish driven by transgenic MYC: Implications for oncogenesis and lymphopoiesis. Leukemia 33: 333–347.

30. Liu, X. J., Y. S. Li, S. A. Shinton, J. Rhodes, L. J. Tang, H. Feng, C. A. Jette, A. T. Look, K. Hayakawa, and R. R. Hardy. 2017. Zebrafish B cell development without a pre-B cell stage, revealed by *cd79* fluorescence reporter transgenes. J Immunol 199: 1706–1715.

31. Langenau, D. M., H. Feng, S. Berghmans, J. P. Kanki, J. L. Kutok, and A. T. Look. 2005. Cre/lox-regulated transgenic zebrafish model with conditional myc-induced T cell acute lymphoblastic leukemia. Proc Natl Acad Sci USA 102: 6068–6073.

32. Bankhead, P., M. B. Loughrey, J. A. Fernandez, Y. Dombrowski, D. G. McArt, P. D. Dunne, S. McQuaid, R. T. Gray, L. J. Murray, H. G. Coleman, J. A. James, M. Salto-Tellez, and P. W. Hamilton. 2017. QuPath: Open source software for digital pathology image analysis. Sci Rep 7: 16878.

33. Schindelin, J., I. Arganda-Carreras, E. Frise, V. Kaynig, M. Longair, T. Pietzsch, S. Preibisch, C. Rueden, S. Saalfeld, B. Schmid, J. Y. Tinevez, D. J. White, V. Hartenstein, K. Eliceiri, P. Tomancak, and A. Cardona. 2012. Fiji: An open-source platform for biological-image analysis. Nat Methods 9: 676–682.

34. Cardona, A., S. Saalfeld, J. Schindelin, I. Arganda-Carreras, S. Preibisch, M. Longair, P. Tomancak, V. Hartenstein, and R. J. Douglas. 2012. TrakEM2 software for neural circuit reconstruction. PLoS One 7: e38011.

35. Cignoni, P., E. Gobbetti, R. Pintus, and R. Scopigno. 2008. Color Enhancement for Rapid Prototyping. In VAST 2008: The 9th International Symposium on Virtual Reality, Archaeology and Intelligent Cultural Heritage, Braga, Portugal. 9-16.

36. Andrews, S., J. Gilley, and M. P. Coleman. 2010. Difference tracker: ImageJ plugins for fully automated analysis of multiple axonal transport parameters. J Neurosci Methods 193: 281–287.

37. Bushnell, B. 2014. BBMap: A fast, accurate, splice-aware aligner. In Conference: 9th Annual Genomics of Energy & Environment Meeting, Walnut Creek, CA, March 17-20, 2014, United States. Medium: ED.

38. Okonechnikov, K., A. Conesa, and F. García-Alcalde. 2016. Qualimap 2: Advanced multi-sample quality control for high-throughput sequencing data. Bioinformatics 32: 292–294.

39. Liao, Y., G. K. Smyth, and W. Shi. 2019. The R package Rsubread is easier, faster, cheaper and better for alignment and quantification of RNA sequencing reads. Nucleic Acids Res 47: e47.

40. Love, M. I., W. Huber, and S. Anders. 2014. Moderated estimation of fold change and dispersion for RNA-seq data with DESeq2. Genome Biol 15: 550.

41. Stephens, M. 2016. False discovery rates: A new deal. Biostatistics 18: 275–294.

42. Smith, A. C. H., A. R. Raimondi, C. D. Salthouse, M. S. Ignatius, J. S. Blackburn, I. V. Mizgirev, N. Y. Storer, J. L. O. de Jong, A. T. Chen, Y. Zhou, S. Revskoy, L. I. Zon, and D. M. Langenau. 2010. High-throughput cell transplantation establishes that tumor-initiating cells are abundant in zebrafish T cell acute lymphoblastic leukemia. Blood 115: 3296–3303.

43. Park, G., J. Burroughs-Garcia, C. A. Foster, A. Hasan, C. Borga, and J. K. Frazer. 2020. Zebrafish B cell acute lymphoblastic leukemia: new findings in an old model. Oncotarget 11: 1292–1305.

44. Traver, D., B. H. Paw, K. D. Poss, W. T. Penberthy, S. Lin, and L. I. Zon. 2003. Transplantation and in vivo imaging of multilineage engraftment in zebrafish bloodless mutants. Nat Immunol 4: 1238–1246.

45. Liang, Z., X. Dong, Z. Zhang, Q. Zhang, and Y. Zhao. 2022. Age-related thymic involution: Mechanisms and functional impact. Aging Cell 21: e13671.

46. Castro, R., D. Bernard, M. P. Lefranc, A. Six, A. Benmansour, and P. Boudinot. 2011. T cell diversity and TCR repertoires in teleost fish. Fish Shellfish Immunol 31: 644–654.

47. Koch, U., and F. Radtke. 2011. Mechanisms of T cell development and transformation. Annu Rev Cell Dev Biol 27: 539–562.

48. Zhang, Y. A., I. Salinas, J. Li, D. Parra, S. Bjork, Z. Xu, S. E. LaPatra, J. Bartholomew, and J. O. Sunyer. 2010. IgT, a primitive immunoglobulin class specialized in mucosal immunity. Nat Immunol 11: 827–835.

49. Fillatreau, S., A. Six, S. Magadan, R. Castro, J. O. Sunyer, and P. Boudinot. 2013. The astonishing diversity of Ig classes and B cell repertoires in teleost fish. Front Immunol 4: 28.

50. Zhang, Y., and D. L. Wiest. 2016. Using the zebrafish model to study T cell development. Methods Mol Biol 1323: 273–292.

51. Rezzani, R., L. Nardo, G. Favero, M. Peroni, and L. F. Rodella. 2014. Thymus and aging: Morphological, radiological, and functional overview. Age (Dordr) 36: 313–351.

52. Borga, C., C. A. Foster, S. Iyer, S. P. Garcia, D. M. Langenau, and J. K. Frazer. 2019. Molecularly distinct models of zebrafish MYC-induced B cell leukemia. Leukemia 33: 559–562.

