## Supplementary material for "Dynamic Changes in Lymphocyte Populations Establish Zebrafish as a Thymic Involution Model": SF 1-3

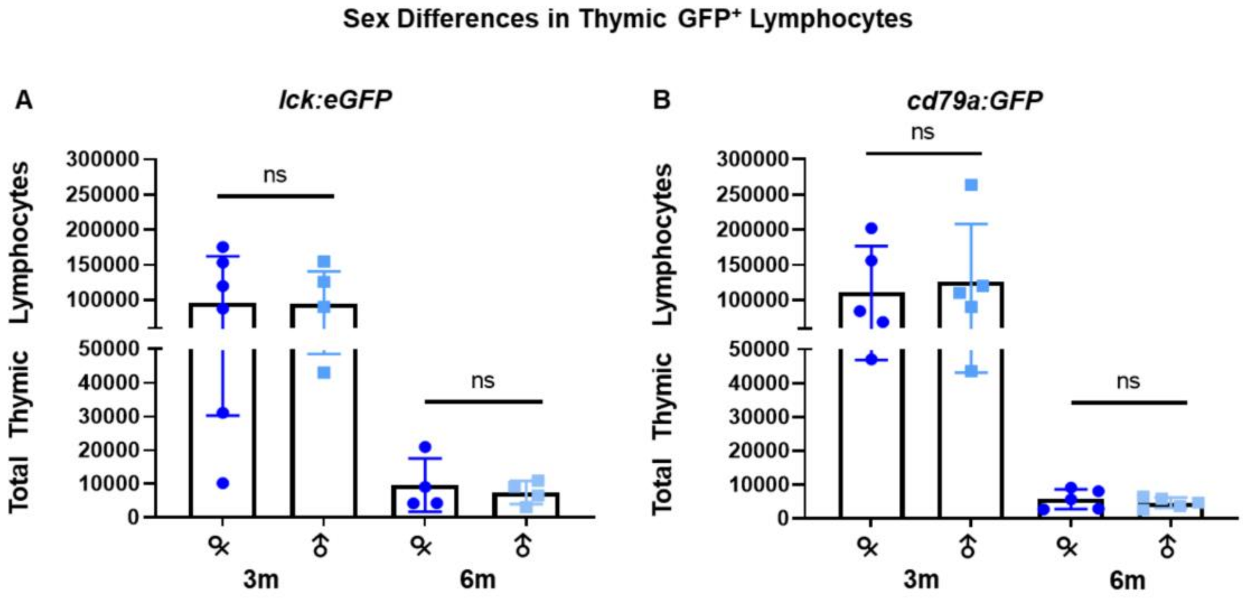


**Supplemental Figure 1: Sex differences in thymic GFP⁺ lymphocytes.** Total thymocytes from (**A**) *lck:eGFP* fish at 3m (F, n=6; M, n=5) and 6m (F, n=4; M, n=4) and (**B**) *cd79a:GFP* fish at 3m (F, n=5; M, n=5) and 6m (F, n=5; M, n=5); ns = not significant *p*-values by 2-way ANOVA tests.

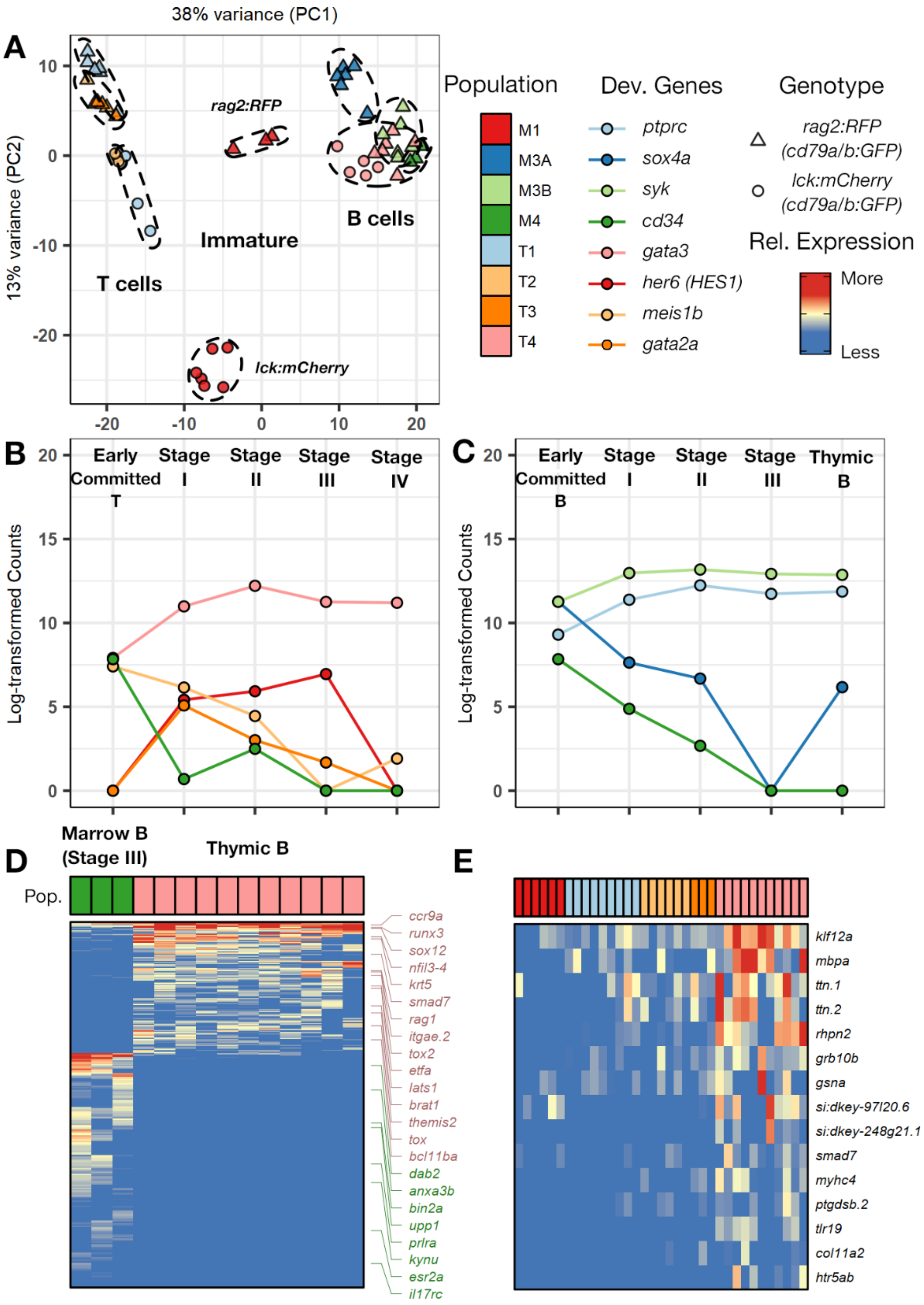

A
38% variance (PC1)
13% variance (PC2)
rag2:RFP
lck:mCherry
T cells
Immature
B cells
Population
M1
M3A
M3B
M4
T1
T2
T3
T4
Dev. Genes
ptprc
sox4a
syk
cd34
gata3
her6 (HES1)
meis1b
gata2a
Genotype
rag2:RFP (cd79a/b:GFP)
lck:mCherry (cd79a/b:GFP)
Rel. Expression
More
Less
B
Early Committed T
Stage I
Stage II
Stage III
Stage IV
Log-transformed Counts
C
Early Committed B
Stage I
Stage II
Stage III
Thymic B
Log-transformed Counts
D
Marrow B (Stage III)
Thymic B
Pop.
ccr9a
runx3
sox12
nfil3-4
krt5
smad7
rag1
itgae.2
tox2
etfa
lats1
brat1
themis2
tox
bcl11ba
dab2
anxa3b
bin2a
upp1
prlra
kynu
esr2a
il17rc
E
klf12a
mbpa
ttn.1
ttn.2
rhpn2
grb10b
gsna
si:dkey-97l20.6
si:dkey-248g21.1
smad7
myhc4
ptgdsb.2
tlr19
col11a2
htr5ab

**Supplemental Figure 2: Stage Specific B and T Cell genes.**

(**A**) Principal component analysis (PCA) of all sequenced samples (n=53) based on the 1000 most-variable genes. T-lineage groups (n=24) cluster at left and B-lineage groups (n=29) cluster in the upper right quadrant. Sample shapes and colors match those of the genotypes and flow cytometric populations in Figures 7-8. (**B**) Normalized counts for select T-lymphopoietic genes across each T cell maturation stage. Early-committed T (*lck:mCherry* M1, red circles in **A**; n=6), Stage I (*rag2*:*RFP*;*cd79a*:*GFP* T2 and T3, light- and dark-orange triangles in **A**; n=6), Stage II (*rag2*:*RFP*;*cd79a* or *cd79b*:*GFP* T1, light-blue triangles in **A**; n=6), Stage III (*lck:mCherry*;*cd79a*:*GFP* T1, light-blue circles in **A**; n=3), and Stage IV (*lck:mCherry*;*cd79a*:*GFP* T2, light-orange circles in **A**; n=3). (**C**) Normalized counts for select B-lymphopoietic genes across each B cell maturation stage, including thymic B cells. Early-committed B (*rag2*:*RFP* M1, red triangles in **A**; n=3), Stage I (*rag2*:*RFP*;*cd79a*:*GFP* or *cd79a*:*GFP* M3A, blue triangles in **A**; n=6), Stage II (*rag2*:*RFP*;*cd79a*:*GFP* or *cd79a*:*GFP* M3B, light-green triangles in **A**; n=6), Stage III (*rag2*:*RFP*;*cd79b*:*GFP* M4, dark-green triangles in **A**; n=3), and thymic B cells (all four genotypes' T4, pink circles and triangles in **A**; n=11). (**D**) Differentially-expressed genes (n=312; 112 up- and 200 down-regulated) in thymic B (T4) vs. stage III marrow B (M4) cells (adjusted *p*-value <0.05, absolute fold-change >1.5, normalized gene count $\geq$10 in at least 2/3 of each group). Select gene abbreviations listed (entire genelist, Supplemental Table 4). (**E**) Up-regulated genes (n=13) in thymic B cells relative to every T cell group and also every marrow B subset.

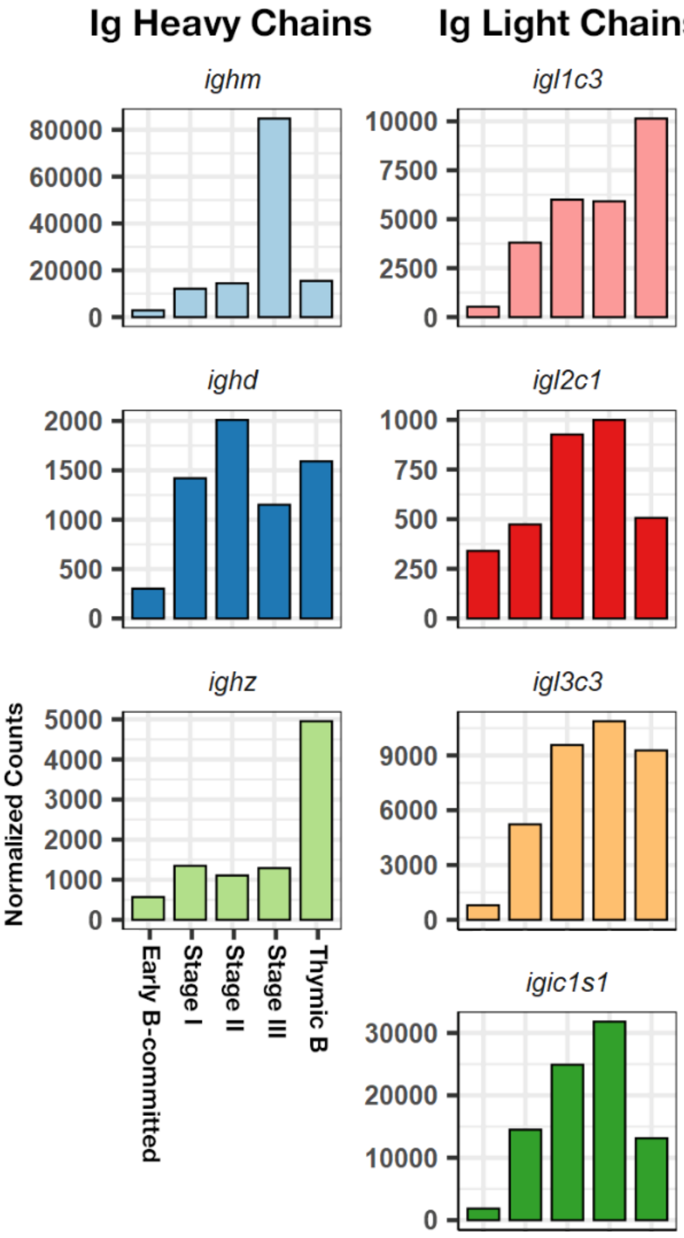


**Supplemental Figure 3: Immunoglobulin Expression by Distinct B Cell Subsets.** Histograms depicting normalized counts of Ig heavy (left column) and light (right column) chain C regions genes at each B cell developmental stage. Counts reflect averages for each group: Early-committed B (n=3), Stage I marrow B (n=6), Stage II marrow B (n=6), Stage III marrow B (n=3), and Thymic B (n=11). Relevant *p*-values are cited in main text.
